# A scRNA-seq Reference Contrasting Living and Early Post-Mortem Human Retina Across Diverse Donor States

**DOI:** 10.1101/2024.09.13.612968

**Authors:** Luning Yang, Yiwen Tao, Qi Pan, Tengda Cai, Yunyan Ye, Jianhui Liu, Yang Zhou, Yongqing Shao, Quanyong Yi, Zen Huat Lu, Lie Chen, Gareth McKay, Richard Rankin, Weihua Meng

## Abstract

**Background:** Current human retina studies predominantly utilize post-mortem tissue, and the sample accessibility constraints make the characterization of the living human retina at single-cell resolution a challenge. Although single-nucleus RNA-seq expands the utility of frozen samples, it provides a nuclear-centric view, potentially missing key cytoplasmic information and transient biological processes. Thus, it is important to generate resources directly from living human retinal tissue to complement existing datasets.

**Methods:** We profiled 106,829 single cells from nine unfrozen human retina samples. Living samples were collected withinLJ10LJmin of therapeutic enucleation and four postmortem samples were collected within 6 h. After standardized dissociation, single-cell transcriptomes were generated using 10x Genomics 3’ RNA-seq and applied scVI to generate batch-corrected integrated atlas. Major cell types and subtypes were annotated through iterative Leiden clustering, canonical markers. Subsequent analyses included differential expression comparisons between cell states and regulon activity profiling to further characterize cellular identities and regulatory networks. Transcriptional dynamics were assessed using RNA velocity, and cell-cell signaling pathways were inferred with CellChat. Key findings were validated in independent samples from two additional donors (four samples) using the identical workflow.

**Results:** We contribute to establishing a reference for retinal cell type proportions and cellular states. Our analysis revealed ELF1-mlCone, a distinct cluster of mlCone photoreceptors identified by distinct transcriptional features. The presence and transcriptional features of this cluster were validated in independent samples. Additionally, by comparing living and post-mortem samples, our study highlights differences in transcriptional dynamics: living tissue preserved coherent RNA velocity streams, enabling clear dynamic state transitions, while post-mortem tissue exhibited disorganized patterns. These findings suggest that using living tissue can improve the capture of active cellular states and transitions.

**Conclusions:** Our atlas provides a single-cell reference contrasting living versus early post-mortem human retina, integrating cell type composition, transcriptional diversity, and functional insights. It may serve as a useful resource for retinal research and for understanding aspects of human retinal biology, particularly given its inclusion of living tissue and diverse pathological states.

## Background

The human retina is a complex and vital component of the eye that plays a crucial role in light capture and the initiation of the vision process [1]. As a highly specialized neural tissue lining the back of the eye, the human retina detects and converts incoming light into electrical signals, forming the foundation of visual perception and interpretation of our surroundings [2]. In general, the retina consists of photoreceptors (rod and cone), signaling neurons (bipolar, horizontal, amacrine, and retinal ganglion cells), and supporting cells (Müller glial cell, microglia, astrocyte) [3]. Each cell type differs in its morphology, functionality, and subtype composition [4]. The retina’s complex structure and function enable humans to process visual information from our environment [5]. However, many questions remain of the structure and function of the human retina [3]. The complex composition of the human retina also makes it vulnerable to various diseases that can significantly impact vision and quality of life, highlighting the importance for improved understanding of the retinal structure and function for the development of effective treatments and preventive measures [6].

Researchers have adopted advanced sequencing technologies to gain deeper insights into retinal biology and disease. Pioneering work by Macosko et al. first applied scRNA-seq to mouse retina [7], later extended to human retina [8]. Single-cell RNA sequencing (scRNA-seq) enables efficient identification of cell type-specific expression patterns, analysis of disease progression stages within the same sample, and exploration of cellular heterogeneity in both physiological and pathological states. For example, Lukowski et al. identified 18 distinct clusters of retinal cells using scRNA-seq, significantly contributing to our understanding of retinal cell diversity [9]. In 2020, Farjood et al. uncovered subpopulations of retinal pigment epithelium cells with stem cell-like properties [10], and Voigt et al. described distinct glial populations in degenerating retinas [11]. Collectively, these studies underscore the utility of scRNA-seq in unraveling the cellular and molecular landscapes of the human retina. Moreover, scRNA-seq provides high-resolution analysis of retinal diseases such as age-related macular degeneration and diabetic retinopathy [12, 13]. Meanwhile, single-nucleus RNA sequencing (snRNA-seq) has facilitated the study of archived or frozen tissues: Wang et al. explored transcription factor collaborations in retinal cells [14], and Liang et al. integrated transcriptome and chromatin accessibility data to identify over 110 cell types [15].

Compared to scRNA-seq, snRNA-seq can be performed on frozen or archived retinal tissues, which is particularly useful where the availability of unfrozen human retinal tissue is limited [16]. The reliance on frozen samples is largely due to the challenges associated with acquiring, preprocessing, and storing human retinal tissue [14]. These challenges encompass not only technical issues but also ethical considerations, such as the difficulty of obtaining samples from living donors [17, 18]. Importantly, snRNA-seq fails to sufficiently capture RNAs localized in the cytoplasm, which are important for understanding post-transcriptional regulation and cellular processes in the retina [19]. A 2009 study by Botling et al. examined the impact of thawing on RNA integrity and gene expression analysis in frozen tissue [20]. They found that the kinetics of RNA degradation after thawing are likely to be tissue specific. This finding is supported by more recent research, including a 2023 review by Thakral et al., which noted that early postmortem interval (PMI) studies on eye tissue found that the type of tissue samples used for early and late PMI had an impact on the stability of RNA post-mortem [21]. This leaves a gap in the field of human retinal research: the lack of resources for scRNA-seq analysis of human retinal samples from living donors, particularly given the fact that living tissue better reflects the *in vivo* cellular transcriptional status more accurately than postmortem or cultured samples [22].

scRNA-seq offers key advantages for retinal studies, capturing both nuclear and cytoplasmic RNAs and enabling the exploration of functional responses in live cells. This technique allows for the analysis of freshly isolated cells while maintaining their viability and function, providing crucial insights into the dynamics of cellular processes such as metabolic activity and stress responses [23].

Unlike many previous studies that focus exclusively on specific regions of the retina, such as the fovea or peripheral retina, our study analyzes the entire adult retina across a larger sample size. While earlier studies have utilized whole retinas, these were limited to a total sample size of three donors [9], making our comprehensive dataset a significant advancement in retinal research. This comprehensive approach ensures that retinal cell types and subtypes are well captured, contributing to a more complete understanding of retinal cellular diversity. By analyzing whole retinas from both living and post-mortem donors, we overcome potential biases introduced by regional sampling and offer a broader perspective on retinal cell heterogeneity. This methodological novelty sets our study apart from earlier scRNA-seq analyses, which often rely on limited retinal regions.

It is worth noting that while previous studies have created retinal atlases and some have used unfrozen postmortem tissue samples, our study is among the first to conduct studies of using living human retinal tissue. By directly comparing living and postmortem samples, we provide additional insights into the advantages of using living tissue in retinal research, building upon and extending previous work in this field.

## Methods

### Retinal tissues from living human donor eyes and cadaveric donor eyes

Human retinal tissues were collected from the Lihuili Hospital (affiliated with Ningbo University) and the Ningbo Eye Hospital (affiliated with Wenzhou Medical University) from June 1, 2023, to July 2, 2024. Study ethics was approved separately by the ethics committees of the Lihuili Hospital, the Ningbo Eye Hospital, and the University of Nottingham, Ningbo, China. Two types of unfrozen full-thickness whole human retinas were collected: (1) living donor samples processed within 10 minutes of surgical enucleation, and (2) post-mortem samples processed within 6 hours of death under temperature-controlled conditions. Our protocol involved dissecting and processing entire retinas to ensure comprehensive coverage of all retinal regions.

Retinal samples were collected following written informed consent from two patients attending the Lihuili Hospital following enucleation due to medical conditions, and eleven samples were collected from the Ningbo Eye Hospital. Of the eleven samples collected from the Ningbo Eye Hospital, four pairs of eyes were collected from recently deceased cadavers, and three from patients following enucleation for medical reasons. Detailed information of the nine samples is shown in Table 1. Donors 01-07 were used as discovery datasets, and donors 08 and 09 were used to perform replications. The overview design of this study is shown in Figure 1A.

**Fig 1.**
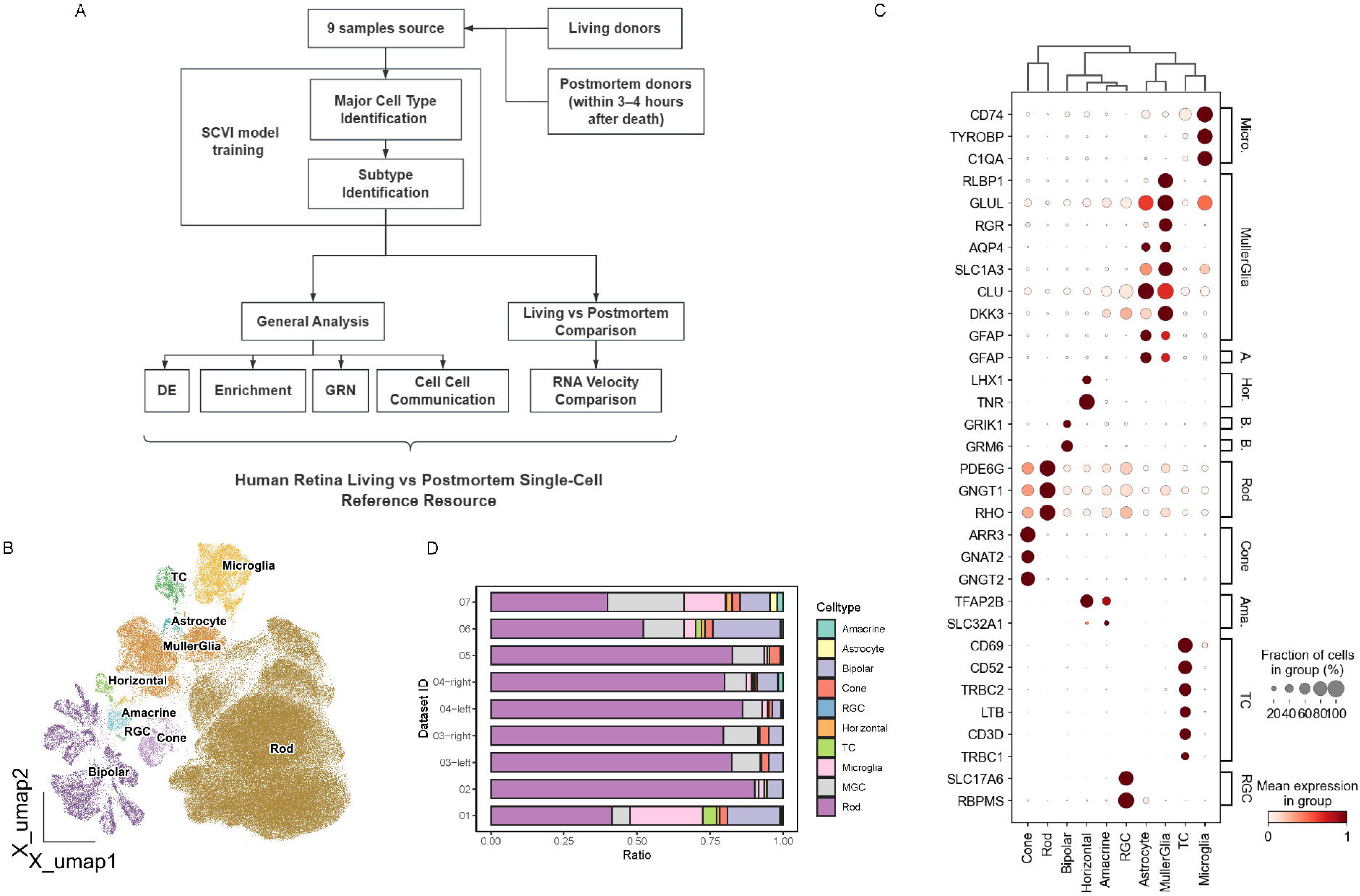
Comprehensive analysis of major cell types in human retinal samples. (A) Overview of the study design workflow for single-cell transcriptomic analysis of human retina. (B) UMAP visualization of integrated human retinal datasets showing distinct clusters of major cell types, including rods, cones, bipolar cells (BC), horizontal cells (HC), amacrine cells (AC), Müller glia cells (MGC), microglia, astrocytes, T cells (TC), and retinal ganglion cells (RGC). (C) Dot plots showing the known markers used to identify major cell types and a selection of genes distinguishing each cluster for retinal samples. Each column corresponds to a specific major cell type, and the rows correspond to a list of key marker genes; brackets on the right of the plot detail the cell type identified by the genes. (D) Stacked bar plot showing the distribution ratios of major cell types across the nine human retina samples.

**Table 1.** Donor clinical and sampling information.

| Donor | Treatment | Eye | Sex | Age | Eye Removal/<br>Death Reason | Diabetic<br>Status | Hospital | PMI |
| --- | --- | --- | --- | --- | --- | --- | --- | --- |
| 01 | living | left | Male | 76 | Corneal staphyloma | DM | L.H. | - |
| 02 | living | left | Male | 55 | Phthisis bulbi | nonDM | L.H. | - |
| 03 | cadaveric | both | Female | 55 | SCT-H-PON | DM | Y.H. | 6 h |
| 04 | cadaveric | both | Male | 55 | BSH | DM+ | Y.H. | 6 h |
| 05 | living | right | Male | 71 | Corneal ulcer | nonDM | Y.H. | - |
| 06 | living | left | Female | 50 | Phthisis bulbi | nonDM | Y.H. | - |
| 07 | living | right | Female | 68 | Corneal penetration | nonDM | Y.H. | - |
| 08 | cadaveric<br>(validation sample) | both | Male | 35 | Traumatic shock | DM+ | Y.H. | 6 h |
| 09 | cadaveric<br>(validation sample) | both | Male | 84 | Prostate malignancy | DM+ | Y.H. | 6 h |

Living: retina samples collected from living humans without freezing. Cadaveric: retina samples collected from recently deceased bodies without freezing. SCT-H-PON:
Severe craniocerebral trauma with hemorrhage in the periphery of the optic nerve. BSH: Brain stem hemorrhage. DM: Samples with diabetes. DM+: Samples with diabetes
and minor retinal hemorrhages. nonDM: Samples without diabetes. L.H.: The Affiliated Li Huili Hospital. Y.H.: The Affiliated Ningbo Eye Hospital of Wenzhou Medical
University. PMI: Estimated postmortem interval (only applicable to cadaveric samples).

### Single-cell RNA sequencing (lab preparation)

Retinal tissue was prepared for scRNA-seq using the Dissociation Kit (LK003150-1bx manufactured by Worthington Biochemical Corporation). Tissue was dissociated, filtered, and treated with red blood cell lysis solution (130-094-183, Miltenyi Biotec). Cell viability was assessed using CountStar. The scRNA-seq libraries were prepared using a total of 10,000 cells per sample. The Chromium Single Cell 3’ Library and Gel Bead Kit V3.1 (10X Genomics, PN1000268) was used to create emulsified single-cell gel beads. Following cell lysis, primers comprising poly-T, barcodes, unique molecular identifiers (UMIs), and a ‘read 1 primer sequence’ in gel bead-in-emulsion (GEM) were used to reverse-transcribe RNA. The addition of sample index and ‘read 2 primer sequence’ was performed through adapter ligation. Following quality control, libraries were 150 bp paired-end nucleotide sequenced using the Illumina NovaSeq platform. Reads were constructed from raw BCL files using Illumina’s bcl2 FASTQ converter. Quality control parameters included: had >3 read sequences; > 20% of bases had a quality rating of < 5; and adapter sequence detected.

### Analysis workflow overview

We processed raw sequencing data using the 10x Genomics Cell Ranger pipeline, followed by downstream analyses in R and Python. Key steps included quality control filtering (DoubletFinder [24]), data integration and batch correction (scVI [25]), differential expression analysis (Seurat [26]), gene ontology enrichment (clusterProfiler [27]), regulon analysis (SCENIC [28]), RNA velocity computation (velocyto [29] and Dynamo [30]), and cell-cell communication inference (CellChat [31]). The specific parameters and versions for each tool are detailed below.

### Generation of single cell data

The 10X Genomics Cell Ranger V8.0.1 pipeline (https://support.10xgenomics.com/single-cell-gene-expression/software/pipelines/latest/what-is-cell-ranger) was used with default parameters to demultiplex and map raw data to the reference genome (GRCh38-2024-A). Alignment, filtering, barcode counting, and UMI (unique molecular identifier) counting were all done by Cell Ranger count on FASTQ files which generates feature barcode matrices using the chromium cellular barcodes, Cell Ranger count, and gene expression algorithms.

### Quality control and doublets removal

For each sample, 10X Genomics Cell Ranger created raw and filtered feature-barcode matrices. Quality control and doublet removal were conducted separately for each dataset to remove and enrich low-quality and enrich high-quality cells, respectively. DoubletFinder (v2.0.4) [24] was used to eliminate doublets. Subsequently, cells of high quality were selected using the Seurat R package (v5.1.0) [26], according to the following quality control criteria: cells with a gene count > 200, UMI count > 400, and a mitochondrial read count proportion < 20% were retained.

### Cell clustering and dimensionality reduction

Scanpy (v1.10.2) [32] was used to analyze the scRNA-seq data. Raw counts were log-normalized, and 5,000 highly variable genes were selected. The scVI model from the scvi-tools package (v0.17.1) [25] was trained using the raw counts of these highly variable genes for dimension reduction and batch correction. Using the 20 resultant latent dimensions, a *k*-nearest neighbors (KNN) graph was created for Leiden clustering and uniform manifold approximation and projection (UMAP) visualization.

Iterative Leiden clustering with parameters of 0.05 to 1.0 was employed to identify cellular partitions within the dataset. In each iteration, a new KNN graph was generated, and Leiden clustering was applied to the scVI batch-corrected latent representation. The calculation of silhouette scores for Leiden clustering at increasing resolutions of 0.1 provided optimal clustering resolution for the scRNA-seq data. The clustering quality for each resolution was assessed using scikit-learn, and the resolution with the highest silhouette score was chosen to identify the most distinct and significant clusters. This was further verified by UMAP plots and cell type specific marker gene expression to ensure the resolution chosen provided the most well-defined and meaningful clusters, enabling the assignment of the first-level annotation of major cell classes. Cells were then divided into smaller groups, and the steps were repeated to finalize subcluster cell type annotations. Subclusters were identified and labeled using known marker genes, differential expression (DE) analysis, and comprehensive enrichment analysis. The optimal cluster resolution for each group was determined by these results and the corresponding silhouette score.

### Comparative analysis of retinal cell type proportions from published studies

To investigate discrepancies in retinal cell type composition across published studies, we performed a systematic reanalysis of publicly available human retinal scRNA-seq datasets. Data was obtained from *Single-cell Atlas of the Human Retina v1.0* on the Human Cell Atlas (HCA) platform (https://data.humancellatlas.org/hca-bio-networks/eye/atlases/retina-v1-0), which integrates major published human retinal studies [33]. The section ‘scRNA-seq of human retina – all cells’ was taken for comparison, composed of five studies [8, 9, 34–36]. Only one study utilized cell-type enrichment (rod depletion and retinal ganglion cells enrichment) for peripheral samples [36]. All other studies profiled unselected whole-retina suspensions. Raw count matrices and associated metadata were downloaded directly from the HCA portal. The datasets represent a compilation of approximately 2.4 million cells from 55 donors across multiple studies. Cell type annotations in these integrated datasets benefit from double annotation, initial annotations by the original study authors followed by harmonization through the HCA curation process, enhancing cross-study comparability and reducing annotation inconsistencies. Retinal regional classifications (foveal centralis, peripheral retina, etc.) were based on the tissue region annotations provided in the HCA metadata, which were originally defined by each contributing study’s sampling methodology. For each dataset, cells were grouped by sample ID, tissue region, and annotated cell type. Cell type proportions were calculated for each sample as the relative fraction of each cell type within that sample’s total captured cells.

### Differential expression (DE) and enrichment analysis

The FindMarkers, FindAllMarkers functions in Seurat were utilized to identify differentially expressed genes (DEGs) across each retinal cell class with the Wilcoxon rank-sum test. The DE results were then used to perform Gene Ontology (GO) enrichment analysis, conducted using ClusterProfiler (v4.10.1) [27] to identify significantly enriched pathways in each population.

### SCENIC and cell-type specificity index analysis

The SCENIC R package was used to identify transcription factor (TF) regulons specific for each cell type using human retinal cones [28]. The raw count gene expression matrix of cones processed with previous quality control steps was used as the input data. After retaining genes with at least three counts and expressed in over one percent of total cells, we used GENIE3 to perform correlation analysis and TF-target correlation assessment. Co-expression modules were identified and refined into regulons based on DNA-motif analysis. Two complementary regulatory region databases were utilized: one encompassing region 10 kb upstream and downstream of transcription start sites, and another focusing on 500 bp upstream and 100 bp downstream, both from the hg38-RefSeq-r80 genome annotation. Regulon activity scores were later calculated using AUCell, followed by binarization to determine the activity state of each regulon in individual cells.

Following the SCENIC analysis, the connection specificity index (CSI) analysis was applied to identify and characterize transcriptional regulatory modules. It was based on regulon activity scores derived from the SCENIC output. CSI quantifies the similarity of regulon activity patterns across cells, enabling the identification of co-regulated gene sets with values ranging from 0 to 1. Higher CSI values indicate greater specificity in co-regulation. We applied a CSI threshold of 0.5 to focus on strongly co-regulated regulons. The resulting CSI matrix was hierarchically clustered using Ward’s method to identify modules of co-regulated genes.

### Dynamo analysis

We used Velocyto [29] to generate spliced and unspliced expression matrices from Cell Ranger output files. These two matrices, along with the raw expression matrix, were imported into Dynamo [30], a python package for RNA velocity analysis. Data preprocessing included normalization and dimension reduction. Gene-specific velocity dynamics were calculated, and RNA velocity-derived metrics were generated, including speed, acceleration, divergence, and ddhodge potential, which quantifies the potential landscape of single-cell transitions by decomposing velocity-derived flows into gradient and curl components, with lower values highlighting likely sink states.

### Cell communication analysis

Cell-cell communication analysis was performed using the CellChat [31] package in R. Data preprocessing steps involved subsetting to signaling genes, identifying overexpressed genes and interactions, and projecting data onto a protein-protein interaction network. Communication probabilities were computed, filtered, and aggregated into a network.

### Validation of ELF1-mlCone

To validate the discovery of the ELF1-mlCone subtype, we performed a replication study using independent retinal samples. Nine samples from donors 01-07 were used to generate the atlas, and four retinal samples from two additional donors (Donors 08 and 09) were utilized for the replication. The replication samples were conducted using the same scRNA-seq and analysis pipeline as the discovery datasets. We applied consistent cell type annotation criteria, focusing particularly on the identification of cone subtypes.

## Results

### Single-cell transcriptomic analysis of human retina from living and cadaveric donors

To investigate the cellular landscape of the human retina at single-cell resolution, we collected fresh retinal tissue samples from five living individuals and two paired samples from recently deceased donors. Following quality control, a total of 106,829 high-quality cells were retained for downstream analysis. Clustering was performed on integrated samples, and the resulting clusters were shown in Figure 1B. Individual cells were identified to the major retinal cell types, annotated based on the known markers reported previously [37], including AC (TFAP2B/SLC32A1), astrocyte (GFAP), cone (ARR3/GNAT2/GNGT2), rod (RHO/PDE6G/GNGT1), Müller glia cell (RLBP1), horizontal cells (LHX1/TNR), BC (NETO1), retinal ganglion cells (SLC17A6/RBPMS), microglia (CD74/TYROBP/C1QA), and T cells (CD69/CD52/CD3D) (Figure 1C).

Major cell types exhibited well-isolated, tight clusters (Figure 1B) with markers showing quite distinct and expected distribution among the cell types. In addition, cluster heterogeneity is visible for some cell classes with multiple types, such as BCs. The ACs, astrocytes, BCs, cones, horizontal cells (HCs), T cells, microglia, Müller glia cells (MGCs), and rods were detected in all the analyzed datasets, and retinal ganglion cells (RGCs) were detected in four donors (3, 4, 6 and 7). For all the datasets, rod photoreceptor was the most abundant major type, followed by MGC and BC in our datasets (Figure 1D and Table 2).

**Table 2.**
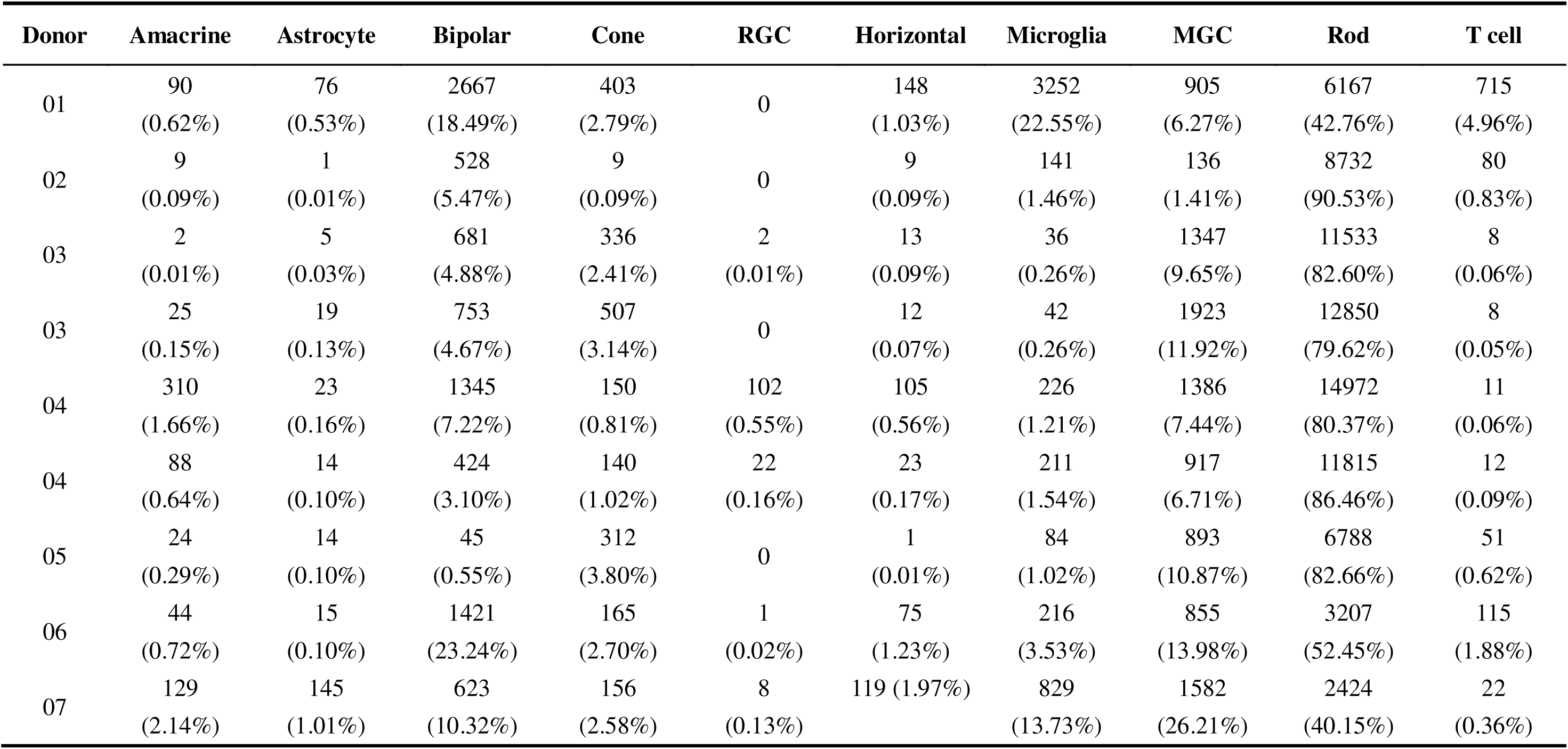
Counts and proportion of major cell types in each human retinal scRNA-seq datasets. The major cell classes include amacrine cells, astrocytes, bipolar cells, cones, retinal ganglion cells (RGCs), horizontal cells, microglia, Müller glia cells (MGCs), rods and T cells.

To validate our results, we compared our data with human retinal scRNA-seq datasets from the HCA platform [33], which integrated five major studies in human retina, as a representative public dataset.

The retinal cell type proportions, categorized by studies and tissue types, were summarized for each sample of published studies (Additional file 2: Table S1), indicating regional variations in cellular composition, but also substantial variation between studies of the same region. This variation likely reflects not only biological differences, but also technical factors such as tissue processing, cell selection, sampling boundaries, and computational annotation. The fovea centralis shows a unique cellular composition, with bipolar cells comprising 29-35% of the population and relatively moderate rod proportions (8-21%). In contrast, the peripheral retina demonstrates markedly different compositions, with rod cells dominating (67 and 79% in Roska/Hafler studies) and reduced bipolar cell percentages except Sanes’ study (22%). These patterns reflect known functional specializations of retinal regions but also raise concerns about the substantial variation in cell type proportions between studies of the same tissue type. Each study reveals distinct technical biases affecting cell type capture, with the Sanes study generally capturing a broader diversity of cell types, the Hafler study showing consistently higher rod proportions, and the Scheetz study demonstrating notably higher MGC capture rates. These differences likely reflect variations in tissue processing methods and cell isolation protocol. At the same time, rare cell types such as HCs, sCone, and specific amacrine subtypes (e.g., GABAergic and glycinergic amacrine cells) are inconsistently captured across datasets.

### Bipolar cells

BCs can be subdivided on the basis of two main distinctions: rod versus cone BCs and ON versus OFF BCs, as shown in Figure 2A. The former is associated with synaptic input, while the latter distinguishes the signaling mechanism in dendrites and the level of axonal stratification [38]. BCs were divided into subtypes previously characterized by common marker genes for each BC subpopulation across all the datasets (Figure 2B). A total of 6 ON BC and 6 OFF BC subclusters were identified in the human datasets (Figure 2C).

**Fig 2.**
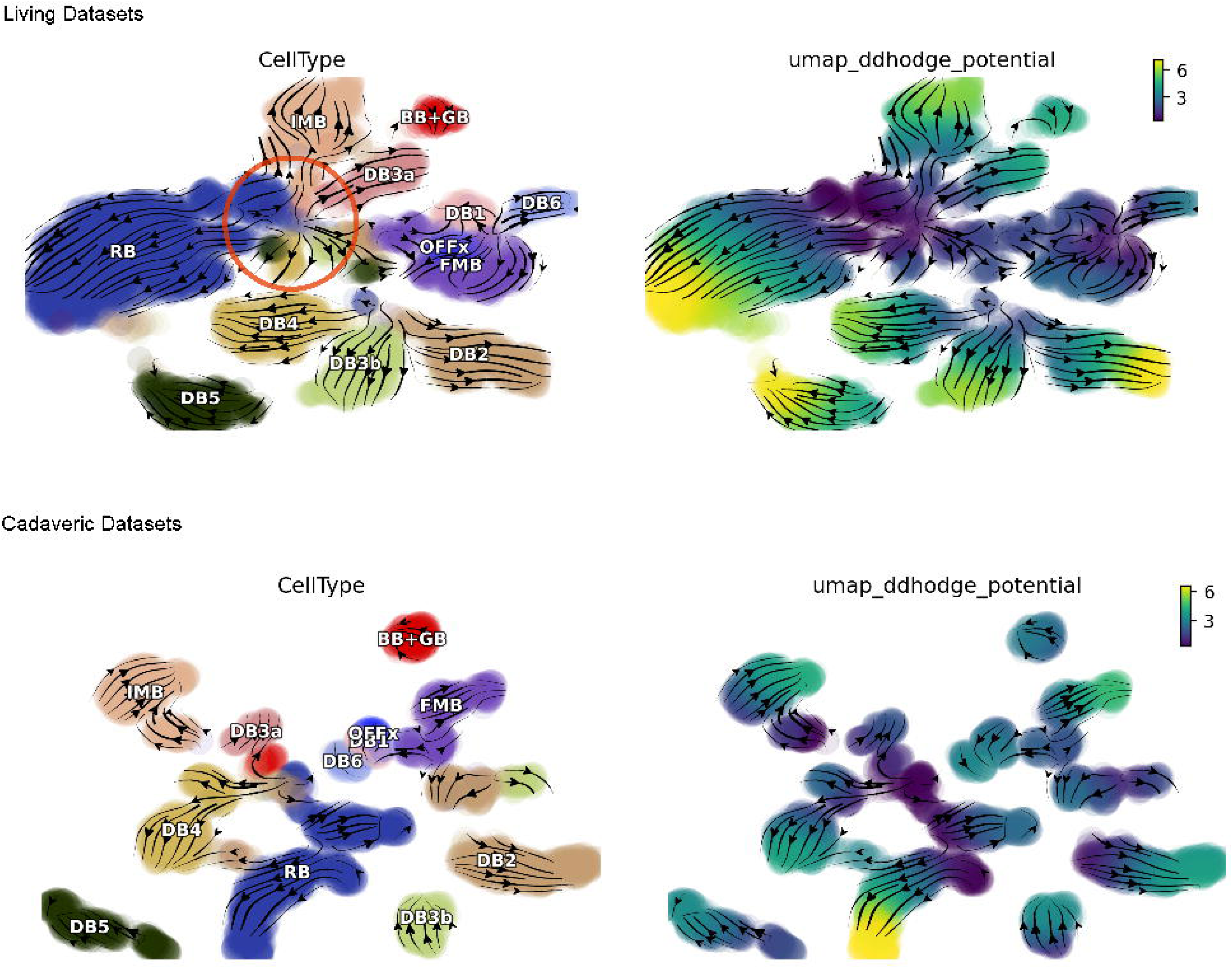
Characterization of bipolar cells (BCs) amacrine cell (AC) and subtypes in the human retina. (A) Illustration of 12 bipolar cell subtypes identified in the human retina. The diagram shows OFF bipolar cells (DB1, DB2, DB3a, DB3b, FMB, OFFx) and ON BCs (RBC, DB4, DB5, DB6, BB+GB, IMB), with distinct morphologies reflecting their light response properties. (B) Dot plots showing the known markers used to identify BC subtypes and a selection of genes distinguishing each BC population. Each row corresponds to a specific BC subtype, and the columns correspond to a list of key marker genes. Brackets on the top of the plot detail the bipolar cell subtype that these genes identify. (C) UMAP visualization of BCs showing 12 distinct populations. (D) Dot plot showing the expression levels of key marker genes in five AC clusters. Rows correspond to each cluster, and the columns correspond to a list of key marker genes, brackets on the top detail the cell type that these genes identify. (E) UMAP visualization of ACs showing five distinct populations: GABA-AC1, GABA-AC2, GABA-AC3, Gly-AC, and Intermediate-AC. (F) The dot plot illustrates the results of GO enrichment analysis for differentially expressed genes in various amacrine cell population. The y-axis displays the top 10 significantly enriched GO terms for each cell subtype, encompassing biological processes, cellular components, and molecular functions. The x-axis represents the different amacrine cell subtypes. Each dot’s size corresponds to the gene count, while the color intensity indicates the gene ratio. The color scale ranges from yellow (lower gene ratio) to purple (higher gene ratio).

The ON BC subtypes, such as DB5, DB6, BB+GB, and IMB, respond to light onset and transmit signals via mGluR6 receptors [39]. OFF BC subtypes, including DB1, DB2, DB3a, DB3b, FMC, and OFFx, hyperpolarize in response to light offset, utilizing ionotropic glutamate receptors (AMPA and Kainate) [40]. Rod bipolar cells (RBCs) are crucial for scotopic (low-light) vision, transmitting signals from rod photoreceptors primarily to AII ACs, which then relay the information to cone BC cells and ultimately to RGCs [41]. This classification highlights the diverse roles and pathways of BCs in visual processing. The DE analysis results between BC subtypes were calculated and provided (Additional file 3: Table S2).

### Amacrine cells

Retinal amacrine cells play a complex role in visual signal processing, forming synapses with multiple cell types, including BCs, RGCs, and other ACs, providing both feedback inhibition to BCs and feedforward inhibition to RGCs, shaping the visual signal. We identified five distinct AC populations based on key markers (Figure 2D-E): three GABAergic clusters (marked by GAD1/GAD2), one glycinergic cluster (high SLC6A9, GlyACs), and an intermediate cluster showing mixed GABA/glycine characteristics. The three GABAergic populations showed distinct signatures characterized by high expression of MEIS2/BASP1 (GABA-AC1), SOX5/PCP4 (GABA-AC2), and CCDC39/RBFOX1 (GABA-AC3). The DE results of each AC cluster were provided in Additional file 4: Table S3.

To further verify the function of the three different GABAergic ACs, we performed GO enrichment analyses for each cluster (Figure 2F, Additional file 1: FigureS1). Except the GlyACs and intermediate ACs, the results of pathway analysis of these three different GABAergic ACs demonstrate that they have distinctly different functions.

GABA-AC1 is particularly enriched in pathways related to “glutamatergic synapses” and “synaptic membranes,” suggesting involvement in a range of synaptic interactions, potentially encompassing both inhibitory (GABAergic) and excitatory (glutamatergic) signals. Pathways such as “regulation of trans-synaptic signaling” and “modulation of chemical synaptic transmission” underscore a possible role in fine-tuning neurotransmitter release. References to “distal axons” and “growth cones” in the enrichment analysis further suggest that GABA-AC1 may be adapted for active synaptic interactions.

Moreover, GABA-AC1 also exhibits the highest association with developmental processes among the GABA-ACs.

GABA-AC2 is enriched in pathways involving “postsynaptic density” and “postsynaptic specialization,” which could indicate functions in refining synaptic inputs. Its links to photoreceptor outer segments and various ciliary structures suggest possible involvement in retinal growth or maintenance. In contrast, GABA-AC3 shows a marked focus on ion channel activity and calcium signaling, particularly voltage-gated and monoatomic ion channels, pointing to a potential role in rapid signal modulation. Enrichment in pathways related to synapse organization and regulation of trans-synaptic signaling further implies that GABA-AC3 may influence broader synaptic communication. Taken together, these findings support the idea that each GABA-AC cluster contributes in a specialized way to inhibitory processing in the retina, highlighting the functional diversity of amacrine cells.

### Cone photoreceptors

We identified three cone clusters that correspond to distinct cell states, referred to as mlCone, sCone, ELF1-mlCone (Figure 3A). The main populations mlCone (medium/long-wavelength sensitive) and sCone (short-wavelength sensitive) were annotated by their expression of opsins OPN1LW/MW and OPN1SW respectively (Figure 3B). In addition to the three cone subpopulations, we detected a small cluster labeled “CR.” This cluster lacks canonical cone markers and shows distinct gene expression patterns. The CR cluster was predominantly present in sample 05 (corneal ulcer sample, 80.45% of cones), may reflect cornea-derived or repair-like cells instead of a genuine cone subtype (Figure 3C). We retain this cluster for completeness but do not interpret it as a novel retinal cell population.

**Fig 3.**
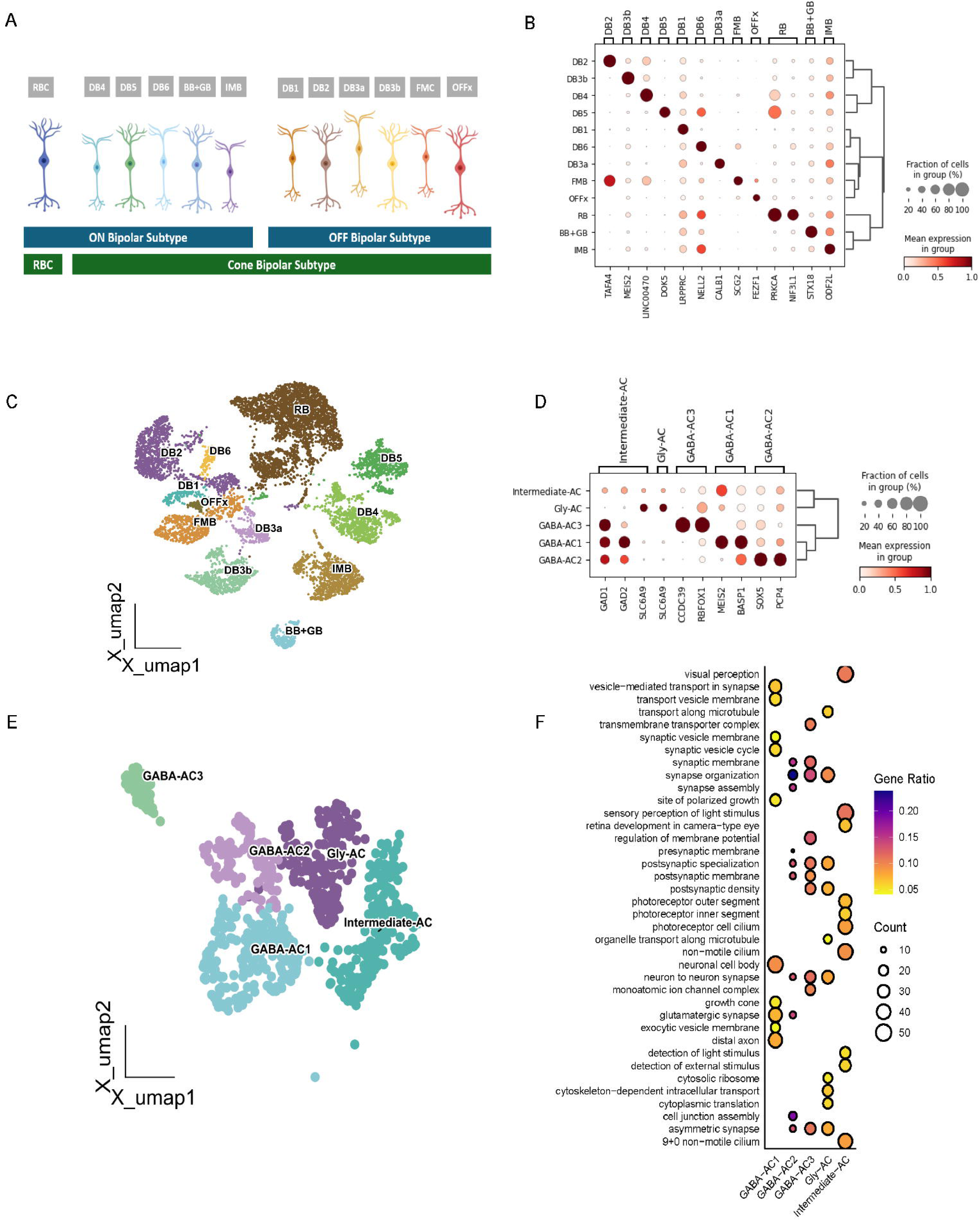
Characterization of cone subclusters in the human retina. (A) UMAP visualization of cone photoreceptor populations showing four populations: sCone, ELF1-mlCone, mlCone and CR cluster. (B) Dot plot showing markers used to identify each cone subpopulation. Each row corresponds to a specific cone population, and the columns correspond to a list of key markers, brackets on the top of the plot detail the cluster that these genes identify. The size of the dots represents the fraction of cells in each group, and the color intensity represents the mean expression level. (C) Stacked bar plot showing the distribution of cone subpopulations within each sample. (D) Dot plots showing markers used to differentiate specifically between mlCone and ELF1-mlCone. (E) GO enrichment analysis comparing ELF1-mlCone vs mlCone populations. Bar plot shows significantly enriched pathways, with red bars indicating upregulated and blue bars showing downregulated pathways in ELF1-mlCone compared to mlCone. (F) RSS (Regulon Specificity Score) ranking analysis showing transcription factor activities in ELF1-mlCone population. SCENIC analysis was performed to identify cell-type-specific transcription factor regulons. The x-axis represents the rank of each regulon, and the y-axis shows the RSS scores, which quantify the specificity of each regulon to the ELF1-mlCone population (higher scores indicate greater specificity). ELF1 shows the highest specificity score, indicating its distinctive association with this cone subpopulation. (G) Connection Specificity Index (CSI) correlation heatmap of regulon modules in cone photoreceptors. CSI analysis was applied to SCENIC-derived regulon activity scores to identify co-regulated transcriptional modules. The heatmap shows correlation patterns between regulons, with color intensity representing correlation strength (dark blue: high positive correlation, light colors: low correlation). Four distinct modules are identified through hierarchical clustering and numbered (Module 1-4). Key regulons with the highest RSS rank values in ELF1-mlCone identified by SCENIC analysis are labeled on the right side, showing their modular organization and co-regulation patterns.

We also revealed a cluster of mlCone photoreceptors exhibiting specialized transcriptional features, which we termed ELF1-mlCone, characterized by enhanced expression of synaptic machinery genes including KCNB2 (voltage-gated K+ channel), CADPS, and RIMS2 (Figure 3D). While maintaining core cone markers, ELF1-mlCones showed coordinated upregulation of synaptic components compared to traditional mlCones, suggesting functional adaptation rather than a distinct cell type.

To validate that ELF1-mlCone represents a distinct biological entity rather than technical artifacts (such as doublets), we performed comprehensive analyses. ELF1-mlCone existed consistently across samples. As shown in Additional file 5: Table S4, the ratios of ELF1-mlCone over the total size of mlCones varied between 1-4%. The highest ratio is 3.9% and there is a sample that shows 0 but should be excluded due to insufficient total cell numbers (only 9 cones). Excluding the invalid sample, the ratios consistently show that approximately 1.5-2.5% of mlCones were of ELF1-mlCone state. The consistent ratio of ELF1-mlCone to mlCone populations across samples suggests that the ELF1-mlCone cluster reflects a specific cellular characteristic or response within mlCones. Moreover, the violin plots show consistent expressed gene counts and total gene expression distributions across cones and bipolar cells, including ELF1-mlCones (Additional file 1: Figure S2A). It demonstrated distinct expression patterns rather than mixed signatures typical of doublets. If ELF1-mlCones were doublets, they would show approximately doubled gene and transcript counts compared to other populations. Additionally, the mitochondrial proportion in ELF1-mlCones aligns closely with other cone populations, indicating intact single cells rather than cell aggregates or technical artifacts.

The expression patterns in Additional file 1: Figure S2B further supported this conclusion. ELF1-mlCones maintain expression of cone-specific markers while showing no significant expression of BC markers. Key synaptic genes like KCNB2, CADPS, and RIMS2 show coordinated enhancement in ELF1-mlCones compared to mlCones but maintain similar relative expression patterns. This suggested a refinement of existing cone machinery rather than the mixed expression signature that would be expected from a doublet of cone and cone-bipolars. Notably, bipolar cell-specific markers (PRKCA, SG2, FEZ1) are absent in ELF1-mlCones, while cone markers like ARR3 are maintained. The systematic nature of these expression changes, with coordinated enhancement of specific pathways rather than random mixing of cell type signatures, strongly distinguish ELF1-mlCone from the cone bipolar cells.

From the DE analysis results between mlCone and ELF1-mlCone, the most significantly upregulated genes in ELF1-mlCone include KCNB2 (log2FC = 1.55), EYS (log2FC = 1.33), and CADPS (log2FC = 1.48) as shown in Additional file 6: Table S5. These genes are involved in synaptic transmission and structural organization, with remarkably small percentage differences in expression between populations. This pattern suggests a modulation of existing pathways rather than activation of entirely new ones. The consistent expression of these genes across both populations, with enhanced levels in ELF1-mlCone, indicates a refinement of function rather than a fundamental change in cell identity. Notably, the analysis reveals coordinated downregulation of housekeeping genes in ELF1-mlCone, including EEF1A1 (log2FC = –1.85), PRDX2 (log2FC = –2.47), and GAPDH (log2FC = –2.0).

The expression patterns show remarkable continuity between the two populations. Most differentially expressed genes maintain detectable expression levels in both populations, with differences primarily in magnitude rather than presence/absence. This is particularly evident in synaptic genes like RIMS2 (log2FC = 1.10) and DLG2 (log2FC = 1.45), which show enhanced expression in ELF1-mlCone while maintaining substantial levels in mlCone. The small percentage differences in these key functional genes strongly support a model where ELF1-mlCone represents a specialized state rather than a distinct cell type.

The GO enrichment analysis highlighted pathways consistent with specialized functions in the ELF1-mlCone cluster (Figure 3E). The upregulated pathways in ELF1-mlCone are primarily focused on microtubule-based transport, cilium assembly/organization, and protein polymerization, with significant enrichment scores (-log10(qvalue) > 10). These structural modifications align with the enhanced synaptic machinery observed in the differential expression analysis. Notably, the downregulated pathways reveal a coordinated metabolic adaptation, with strong enrichment in cytoplasmic translation (-log10(qvalue) > 50), aerobic respiration, oxidative phosphorylation, and mitochondrial respiratory chain complex assembly. This systematic downregulation of basic cellular processes suggests a metabolic trade-off where ELF1-mlCones reduce energy-intensive housekeeping functions to support enhanced synaptic specialization.

### Key TFs in ELF1-mlCone

On this basis, for further experimentation and validation, we performed SCENIC analysis to profile cell state specificity and TF activity. The top 40 regulons revealed ELF1 as a primary transcription factor, along with other significant regulators including SETDB1, RCOR1, and SREBF1 (Figure 3F, Additional file 7: Table S6).

The regulon analysis identified distinct functional groups of transcription factors. ELF1, the defining transcription factor of this state, showed strong activity along with SETDB1, a histone methyltransferase involved in retinal development and photoreceptor maintenance. RCOR1, a transcriptional co-repressor involved in neuronal gene silencing, and SREBF1, which regulates lipid metabolism, were also highly active. These regulators align with the observed enhancement of synaptic machinery and metabolic adaptation in ELF1-mlCones. CSI analysis revealed four distinct regulon modules, with ELF1-mlCone key transcription factors primarily clustering in modules two and four (Figure 3G, Additional file 8: Table S7). Module two encompassed factors involved in chromatin modification (SETDB1, PHF8), transcriptional regulation (ETV3), and DNA repair (XRCC4), supporting fundamental cellular processes through epigenetic and transcriptional control. Module four contained key regulators including ELF1 and its extended network, metabolic regulators (SREBF1, PRKAA1), and broad transcriptional modifiers (SIN3A, STAT5B), consistent with the metabolic adaptation and specialized state maintenance observed in differential expression analysis. The distinct organization of transcriptional modules (chromatin modifiers in module two, metabolic regulators in module four) suggests a coordinated program where epigenetic regulation maintains cell identity while metabolic adaptation supports enhanced synaptic function.

### Cell to cell interaction inside cones

The cell-cell interaction analysis reveals distinct communication patterns between cone populations (Additional file 9: Table S8). For ELF1-mlCone and mlCone, while both populations maintain core photoreceptor interactions, they show different signaling profiles. mlCones demonstrate broader cellular communication, particularly through NAMPT-INSR signaling with multiple cell types (prob=0.0008-0.0145; CellChat-inferred communication strength) and CNTN1-NRCAM, where mlCones exhibit consistent interactions across neuronal populations (prob=0.0464-0.0493), including bipolar cells (prob=0.0466), amacrine cells (prob=0.0464), and ganglion cells (prob=0.0493). In contrast, ELF1-mlCones exhibit more focused communication patterns. They show enhanced CNTN1-NRCAM signaling with specific targets, particularly with bipolar cells (prob=0.0471) and homotypic interactions (prob=0.0853), while reducing broader cellular interactions. This selectivity is further evidenced by enhanced CDH2-CDH2 homotypic interactions (prob=0.129 vs 0.088 in mlCones) and NEGR1-NEGR1 interactions (prob=0.199 vs 0.120 in mlCones). The interaction networks suggested that ELF1-mlCones maintain selective signaling patterns.

### Replication confirms a transcriptionally distinct ELF1-mlCone cluster

To verify the existence of the ELF1-mlCone, a validation study was conducted using four additional human retinal samples. Figure 4A is the UMAP plot of cone cells, which reveals three clusters corresponding to ELF1-mlCone, mlCone, and sCone. ELF1-mlCone is located as a separate cluster, supporting its identity as a distinct cone subpopulation. Figure 4B is the gene expression dot plot of the key marker of the three cone populations, which was identified in the previous experiments. The heatmap shows high expression of KCNB2 and EYS in the ELF1-mlCone compared to mlCone and sCone. It revealed that ELF1-mlCone is characterized by high expression of KCNB2 and EYS, distinguishing them from mlCones and sCones.

**Fig 4.**
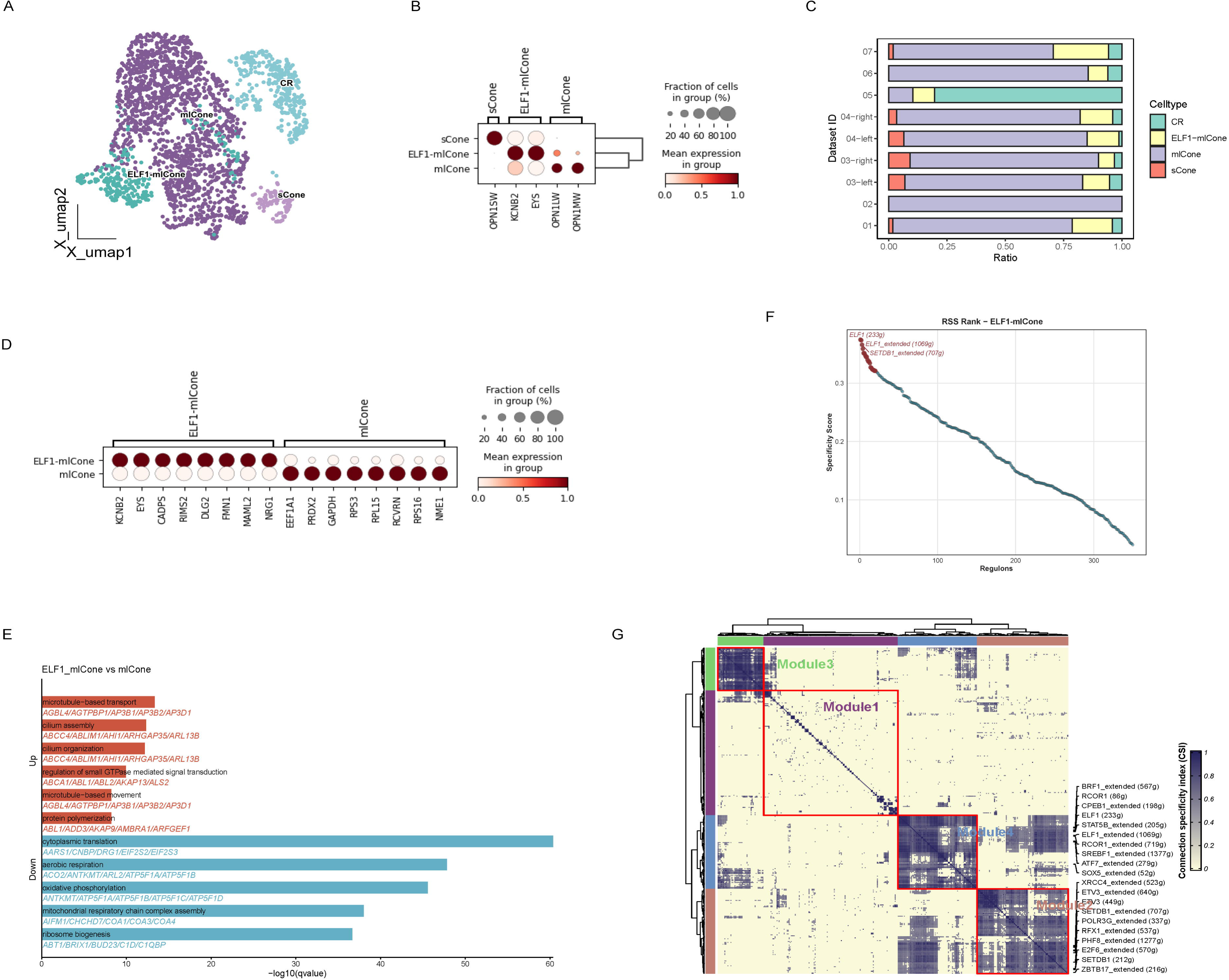
Replication study confirming the existence of ELF1-mlCone and analysis of other cell types. (A) UMAP plot illustrating the clustering of cone based on gene expression profiles, with each color representing a different subpopulation: mlCone (orange), ELF1-mlCone (blue), and sCone (green). (B) Gene expression dot plot of key markers for cone subpopulations. The size of each dot represents the fraction of cells expressing the gene, while the color intensity indicates the mean expression level. Dot plots summarizing the known markers used to identify each cone cell subpopulation. Each row corresponds to a specific cone state, and the columns correspond to a list of key marker genes. Brackets on the top of the plot detail the cluster that these genes identify. The size of the dots represents the fraction of cells in each group, and the color intensity represents the mean expression level. (C) Density plot showing the expression distribution of OPN1LW, OPN1MW, and OPN1SW across the cone populations. The contour lines represent different levels of cell density, with warmer colors indicating higher density. (D) Differential expression analysis across major retinal cell types (Astrocyte, RGC, TC, Microglia, MGC, and Rod). The y-axis shows average log2 fold change, with points colored by significance and expression magnitude: dark gold for highly upregulated genes (log2FC > 1), teal for highly downregulated genes (log2FC < –1), and gray for genes with smaller fold changes. The colored central bar indicates cell type identity. Top 5 up and down regulated genes are labeled for each cell type. (E) Heatmap of hallmark pathway enrichment across retinal cell types using irGSVA. Rows represent MSigDB hallmark gene sets, columns represent cell types (Astrocyte, MGC, Microglia, RGC, Rod, TC). Red indicates pathway upregulation, blue indicates downregulation. Enrichment scores were derived from multiple methods and integrated using Robust Rank Aggregation (* p<0.05, ** p<0.01, *** p<0.001, **** p<0.0001).

Moreover, the density plots of OPN1LW, OPN1MW, and OPN1SW, key markers of mlCone and sCone, were additional evidence for the unique characteristics of ELF1-mlCone as shown in Figure 4C. OPN1LW and OPN1MW showed very similar expression patterns, with high density in the main cluster of mlCone. Importantly, there’s a distinct region with low density for both these markers that corresponds to the ELF1-mlCone population. Density plots demonstrate that ELF1-mlCones occupy a region where traditional cone markers (OPN1LW, OPN1MW, and OPN1SW) show low expression, supporting their distinct transcriptomic profile. These replication results support the identification of ELF1-mlCone as a distinct transcriptional cluster within mlCones with a distinct gene expression signature and clustering behavior, consistently observable across multiple human retinal samples.

### Characterization of Additional Retinal Cell Populations

We also revealed distinct molecular signatures for other retinal cell populations. The human retina contains two horizontal cell subtypes (H1 and H2), identified based on LHX1 and ISL1, respectively (Additional file 1: Figure S3) [42]. Microglia, the resident immune cells essential for retinal homeostasis, exhibited high expression of CD74, TYROBP, and C1QA, with GSVA analysis revealing significant enrichment in inflammatory response pathways, particularly in complement activation and phagocytosis. MGCs demonstrated characteristic expressions of retinoid metabolic process genes and glutamate recycling pathways, highlighting their critical role in visual cycle maintenance and neurotransmitter homeostasis. Astrocytes showed robust pathway enrichment for angiogenesis and epithelial-mesenchymal transition, with particularly strong signatures in coagulation pathway gene, positioning them as key regulators of blood-retina barrier integrity. Astrocytes demonstrated robust pathway enrichment for angiogenesis (VEGFA, ITGAV, NRP1) and vascular regulation (PDGFA, JAG1, THBD), with particularly strong signatures in coagulation and epithelial-mesenchymal transition pathways-uniquely positioning them as key regulators of blood-retina barrier integrity. T cells, expressing CD69, CD52, and CD3D markers, were exclusively detected in our living samples due to their transient presence in healthy retina, showing characteristic adaptive immune pathway signatures. RGCs, though less abundant, exhibited distinctive expressions of RBPM, SNCG, and SLC17A6. GSVA analysis of RGCs revealed enrichment in oxidative-phosphorylation and cholesterol homeostasis, while genes associated with axonogenesis (KIF5A, GAP43, STMN2) and potassium channel activity were also highly expressed, reflecting their specialized function in visual signal transmission to the brain.

### Living vs Post-mortem Analysis Validates Transcriptional Dynamics

Distinct transcriptional dynamics between living and post-mortem samples were observed when performing Dynamo RNA velocity analysis of the cone population. Living samples show clear directional flows from mlCone toward ELF1-mlCone, with coherent velocity streams indicating active transcriptional states. The transition boundary between mlCone and ELF1-mlCone populations exhibits particularly strong streaming patterns. In contrast, post-mortem samples show disrupted velocity patterns with less defined transitions, while maintaining basic cluster identities (Additional file 1: Figure S4)

To validate these findings across retinal cell types, we analyzed BCs using separate UMAP embeddings for living and post-mortem samples. In living samples, velocity streams demonstrate robust directional flows with high coherence across bipolar subtypes. The RBC population exhibits particularly strong streaming patterns with dense, parallel velocity arrows. The transitions between cone bipolar subtypes show organized patterns, with a notable convergence zone (marked by red circle) between IMB, DB3a, and DB4 populations (Figure 5). The umap_ddhodge_potential visualization reveals a gradient pattern where the convergence zone shows the lowest potential (purple), with potential increasing (transitioning through green to yellow) as it extends outward to different BC subtypes, following the direction of the velocity arrows.

**Fig 5.**
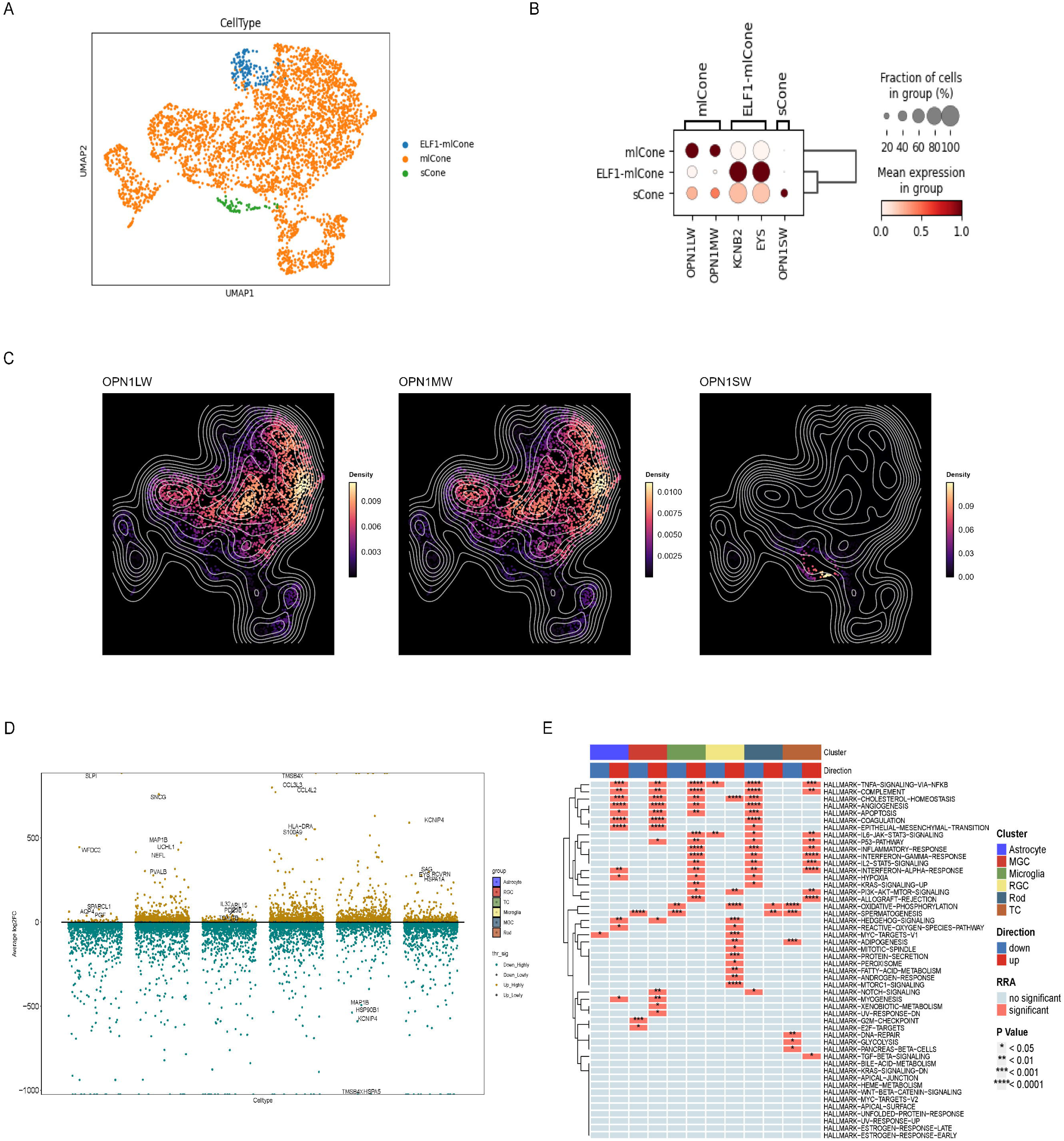
Dynamo RNA velocity analysis of bipolar cell populations in living and cadaveric retinal samples separately. Living Datasets on the top and Cadaveric Datasets at the bottom. Left: UMAP visualization showing distinct bipolar cell subtypes with vector field overlay indicating transcriptional flows. Right: UMAP colored by ddhodge potential field. Vector fields indicate predicted transcriptional flows based on Dynamo analysis, revealing relationships between bipolar cell subtypes and their developmental trajectories.

Cadaveric samples, while preserving basic cluster identities, show markedly altered velocity patterns. The velocity streams become sparse and disorganized across all populations, with particularly pronounced disruptions in the overall connectivity between subtypes. The RBC population maintains its identity but with a distinct breakpoint within itself, where the velocity streams become disrupted and change direction. This breakpoint effectively divides the RBC cluster into two regions with different flow patterns, rather than just showing reduced density or loss of parallel directionality. While The OFF pathway bipolar cells (DB3a, DB3b, DB4) maintain their distinct identities, their spatial organization and connectivity are disrupted. DB3b in particular shows a dramatic shift, appearing more isolated and with reversed velocity vectors compared to its position and flow in living samples. In the ddhodge potential visualization in cadaveric samples, the organized gradient pattern is largely lost, with only the RBC region maintaining a visible yellow-to-purple gradient. The rest of the BC populations show more uniform and muted potential values (predominantly blue-green), lacking the clear directional progression seen in living samples. This suggests that while the RBC population maintains some aspects of its transcriptional dynamics post-mortem, the coordinated potential landscape across other BC subtypes is disrupted.

## Discussion

Our study provides informative single-cell transcriptomic resources derived from human retina, with a particular emphasis on comparing samples from living donors and post-mortem individuals to investigate the impact of tissue viability on scRNA-seq analysis. A key strength of this work lies in the inclusion of retinal tissue from living donors, processed within minutes of enucleation, offering a rare window into cellular transcriptional states that may more closely reflect the in vivo condition compared to studies relying solely on frozen or post-mortem preserved tissue [15]. By incorporating samples from donors with diverse conditions, this dataset also serves as a broad resource for exploring retinal biology across different health statuses, although this heterogeneity also presents limitations.

A primary highlight of this work is the establishment of a reference for the approximate proportions of major retinal cell classes analyzed using consistent whole-retina processing protocols. As noted previously, cell type proportions reported in existing human retinal scRNA-seq datasets vary considerably, potentially due to biological differences (e.g., retinal region sampled) or technical factors like tissue handling and dissociation methods. We attribute the observed differences in cell type distribution across donors to several interconnected technical and biological factors. The pronounced variability in RGC proportions, for instance, is amplified by their inherent low abundance (∼0.2-0.5% of total retinal cells) and their physical fragility, which makes their recovery highly sensitive to minor variations in tissue dissociation. The proportion of rods is also subject to significant variation due to known anatomical differences between the rod-rich peripheral retina and the rod-poor central macula; subtle differences in regional sampling during procurement can therefore lead to large shifts in rod percentages. Finally, due to the nature of proportional data from a fixed cell capture, the relative abundance of cell type like MGC is inversely correlated with the recovery of dominant populations, such as rods, further contributing to inter-sample variability. On the other hand, by performing comparative analysis of region-grouped cell type proportions across major published scRNA-seq studies from HCA, we highlight that methodological differences—including selective cell enrichment, sampling boundaries, dissociation protocols, donor demographics, and analytic pipelines—play a major role in shaping observed cell type proportions in retinal scRNA-seq datasets. These factors must be carefully considered when interpreting cross-study comparisons or using public atlases as reference.

Notably, the true foveal centralis is virtually rod-free, with rods first detected ≈ 300 µm from the pit [43]. Apparent foveal-rod counts in some scRNA-seq studies reflect larger biopsy diameters and pooling strategies rather than biological anomaly. Precise reporting of punch size and eccentricity is essential when comparing atlases.

Beyond characterizing major cell types, we consistently identified a transcriptionally distinct cluster within the mlCone population across multiple samples, which we termed ELF1-mlCone. This cluster is characterized by altered expression of genes related to synaptic machinery (e.g., KCNB2, CADPS, RIMS2) and metabolism (e.g., downregulation of housekeeping genes EEF1A1, PRDX2, GAPDH), along with high activity of the transcription factor ELF1. Cell communication analysis further indicates that while classical mlCones maintain broad interactions with multiple cell types, ELF1-mlCones show more selective interaction patterns, potentially reflecting specialized roles in local synaptic organization. Regarding the biological interpretation of this cluster, different possibilities exist. One is that it represents a specific functional state of mlCone, where the observed molecular features suggest a cellular reprogramming prioritizing synaptic readiness over general metabolic upkeep. Another possibility is that this observed heterogeneity reflects the inclusion of cone cells from different spatial locations within our whole-retina samples. Based on the current scRNA-seq data alone, we cannot definitively distinguish between these possibilities but only verify its consistence using outside datasets. Regardless of its precise identity, identifying and characterizing this reproducibly detected, distinct transcriptomic profile may be useful for future studies requiring precise distinction between cones from different retinal regions, potentially helping to identify regional heterogeneity or ‘cross-contamination’ in samples. Future spatially resolved studies are required to ultimately determine the identity and function of ELF1-mlCone.

A critical aspect of this study was the direct comparison of retinal tissue obtained from living donors and post-mortem samples. Analysis of RNA velocity revealed observable differences in transcriptional dynamic patterns between these two groups across the major cell types examined where analysis was feasible, notably in cones and bipolar cells. Specifically, living samples, processed rapidly after enucleation, generally exhibited more coherent and directional velocity streams, potentially indicative of active transcriptional processes or state transitions. In contrast, post-mortem samples, even those processed within a relatively short interval (6 hours), displayed sparser, more disorganized velocity patterns, despite retaining fundamental cell type identities based on overall gene expression. Postmortem changes, including ATP depletion, ion concentration shifts, enzyme activation, and RNA degradation, all have the possibility to affect cell state, gene expression, and observed developmental trajectories. While previous studies used adult retinal specimens removed of death and dissected within few hours [44], our results indicate that even these relatively short postmortem intervals can lead to significant alterations in cell trajectory. The disruptions observed in postmortem samples may obscure important nuances, particularly when studying subtle cellular transition.

Analyzing tissue processed minutes after enucleation provides an important perspective compared to traditional post-mortem studies. While it is often debated whether trajectory inference or velocity analysis holds biological meaning in fully differentiated tissues like the retina, RNA velocity proved useful beyond development, capturing dynamics in mature tissues. Examples include revealing synaptic changes in Alzheimer’s disease and hippocampal plasticity [45, 46]. Our application of RNA velocity aims not necessarily to reconstruct developmental paths, but to capture and compare the apparent transcriptional dynamics signatures associated with tissue viability. However, factors such as surgical stress, early ischemia and hypoxia, and underlying pathological conditions could potentially influence the dynamics observed in living samples. Although inherent sample heterogeneity further complicates direct comparison, this study, limited by the difficulty of obtaining such rare samples, aims to provide an important perspective. By directly comparing living and post-mortem tissues, our findings offer a helpful reference point, highlighting potential dynamic information loss in commonly used post-mortem datasets and supporting future research aiming to decipher subtle cellular states or responses in the human retina.

## Conclusions

This study provides a single-cell transcriptomic reference resource for the human retina, notably incorporating data from living donors alongside post-mortem samples. We offer a comprehensive baseline for cell type proportions and characterize the transcriptomes of major retinal cell populations. Our comparative analysis reveals differences in measurable transcriptional dynamics (RNA velocity patterns) associated with tissue viability, highlighting the significant impact of sample procurement on such analyses. We also report the consistent detection of a transcriptionally distinct cone cluster, ELF1-mlCone, whose precise identity and functional significance warrant further investigation. We also acknowledge limitations in our study: the transcription factor activity predictions for ELF1-mlCone are based on computational inference rather than direct experimental validation through chromatin immunoprecipitation, proteomic analysis, or functional assays in retinal organoids. Such experimental approaches were beyond the scope of our current study design and resource constraints. However, recognizing these limitations, we employed multiple complementary computational approaches to rigorously characterize this novel cone state in a cautious and systematic manner. This computational validation framework offers evidence to guide future experimental studies to definitively establish the functional significance of the ELF1-mlCone population. To summary, this study serves as a resource for future hypothesis-driven research into human retinal biology, disease mechanisms, and the development of potential therapeutic strategies, highlighting the need to carefully consider tissue source and processing in study design and interpretation.

## Abbreviations

AC: Amacrine cell
BC: Bipolar cell
CSI: Connection Specificity Index
DE: Differentially Expression
DEG: Differentially Expressed Gene
GEM: Gel Bead-in-Emulsion
GO: Gene Ontology
GSVA: Gene Set Variation Analysis
HC: Horizontal cell
HCA: Human Cell Atlas
KNN: k-Nearest Neighbors
MGC: Müller glial cell
mlCone: Medium/long-wavelength-sensitive cone photoreceptor
PMI: Postmortem interval
RBC: Rod bipolar cell
RGC: Retinal ganglion cell
RSS: Regulon Specificity Score
sCone: Short-wavelength-sensitive cone photoreceptor
scRNA-seq: Single-cell RNA sequencing
snRNA-seq: Single-nucleus RNA sequencing
TC: T cell
TF: Transcription factor
UMAP: Uniform Manifold Approximation and Projection
UMI: Unique Molecular Identifier

## Declarations

### Availability of data and materials

The single-cell RNA sequencing data generated in this study have been deposited in Figshare with the DOI: 10.6084/m9.figshare.28227674. Public datasets referenced in the study can be accessed through the *Single-cell Atlas of the Human Retina v1.0* on the Human Cell Atlas platform (https://data.humancellatlas.org/hca-bio-networks/eye/atlases/retina-v1-0).

### Ethics approval and consent to participate

This study was approved by the Ethics Committee of the University of Nottingham Ningbo China; The Lihuili Hospital affiliated with Ningbo University; The Affiliated Ningbo Eye Hospital of Wenzhou Medical University.

## Authors’ information

### Authors and Affiliations

**Nottingham Ningbo China Beacons of Excellence Research and Innovation Institute, University of Nottingham Ningbo China, Ningbo, China, 315100**

Luning Yang, Yiwen Ta, Qi Pan, Tengda Cai, Weihua Meng

**Department of Ophthalmology, Lihuili Hospital affiliated with Ningbo University, Ningbo, China, 315040**

Yunyan Ye

**Department of Cardiology, Lihuili Hospital affiliated with Ningbo University, Ningbo, China, 315040**

Jianhui Liu

**Ningbo Institute of Innovation for Combined Medicine and Engineering, The Affiliated Li Huili Hospital, Ningbo University, Ningbo, China, 315201**

Yang Zhou

**Department of Ophthalmology, The Affiliated Ningbo Eye Hospital of Wenzhou Medical University, Ningbo, China, 315040**

Yongqing Shao

**PAPRSB Institute of Health Sciences, Universiti Brunei Darussalam, Bandar Seri, Begawan, BE1410, Brunei Darussalam**

Zen Huat Lu, Lie Chen

**Centre for Public Health, Institute of Clinical Science, Queen’s University Belfast, Block B, Royal Hospital, Grosvenor Road, Belfast, Northern Ireland, BT12 6BA**

Gareth McKay

**School of Mathematical Sciences, University of Nottingham Ningbo China, Ningbo, China, 315100**

Richard Rankin

## Authors’ contributions

LY (first author) conceived the study with WM, performed all computational analyses, curated the data, prepared the figures and wrote the initial draft. YT, QP and TC assisted with data formatting and quality control, while YY, JL, YZ, YS and QY collected and processed retinal samples. ZL, LC, GM and RR provided critical feedback on the analysis and manuscript. WM (last and corresponding author) secured funding, supervised the project, oversaw methodology and revised the manuscript with input from all authors. WM serves as both the last author and corresponding author for this study.

## Funding

This study was mainly funded by the Pioneer and Leading Goose R&D Program of Zhejiang Province 2023 with reference number 2023C04049 and Ningbo International Collaboration Program 2023 with reference number 2023H025.

## Corresponding authors

Correspondence to Weihua Meng

## Consent for publication

All authors have consent for publication.

## Competing interests

The authors have no conflicts of interest to disclose.

## Additional files

**Additional file 1** (.docx) – Supplementary Figures S1-S4

Word document containing all supplementary figures (S-series) with their accompanying legends.

**Additional file 2** (.xlsx) – Table S1. Cell type proportions across human retinal datasets from multiple studies. This table provides a comprehensive analysis of retinal cell type distributions across different studies (Sanes, Roska, Hafler, Scheetz, and Wong) and tissue regions (fovea centralis, peripheral retina, and chorioretinal region). For each sample, the proportions of major retinal cell types are presented.

**Additional file 3** (.xlsx) – Table S2. Differentially expressed genes among bipolar cell subtypes.

**Additional file 4** (.xlsx) – Table S3. Differentially expressed genes among amacrine cell subpopulations.

**Additional file 5** (.xlsx) – Table S4. Distribution of cone photoreceptor subpopulations across donor samples. This table shows the quantification of cone photoreceptor subtypes (ELF1-mlCone, mlCone, and sCone) across nine donor retinal samples. For each sample, the counts for each population are provided, along with the ratio of ELF1-mlCone to total mlCone population (ELF1-mlCone and mlCone). Samples are labeled by donor ID, with left and right eyes specified where applicable.

**Additional file 6** (.xlsx) – Table S5. Differentially expressed genes of ELF1-mlCone compared to mlCone.

**Additional file 7** (.xlsx) – Table S6. Transcription factor activity ranking in cone photoreceptor subtypes. This table provides RSS (Regulon Specificity Score) rankings of top 40 transcription factors (TFs) in three cone photoreceptor subpopulations (ELF1-mlCone, mlCone, and sCone) separately. For each population, transcription factors are ranked by their RSS scores, with higher scores indicating stronger regulatory activity.

**Additional file 8** (.xlsx) – Table S7. Transcription factor module assignments identified through CSI analysis in cone photoreceptors. This table provides the assignments of transcription factors into four distinct modules (Module 1-4) based on CSI (Connection Specificity Index) analysis. Transcription factors are identified by SCENIC of cone subpopulations.

**Additional file 9** (.xlsx) – Table S8. Cell-cell communication analysis between cone subpopulations and other retinal cells with key pathways include CNTN, CDH, NEGR, and VISFATIN signaling networks.

**Additional file 10** (.xlsx) – Table S9. Differential expression genes of ELF1-mlCone grouped by diabetic status (diabetic versus non-diabetic donors to validate that the metabolic gene expression signature is intrinsic to this cell state rather than disease-related).

## Notes

### Competing Interest Statement

The authors have declared no competing interest.

### Summary of Updates

In this revision, the manuscript title is updated to emphasize the living vs early post-mortem comparison; the newly identified cone population is renamed ELF1-mlCone; the abstract, Results and Discussion are rewritten to centre on tissue-viability effects rather than novelty alone; Figures and legends are revised for clarity; the Methods now include additional comparative-analysis steps

