## Additional file 1 for "A scRNA-seq Reference Contrasting Living and Early Post-Mortem Human Retina Across Diverse Donor States"

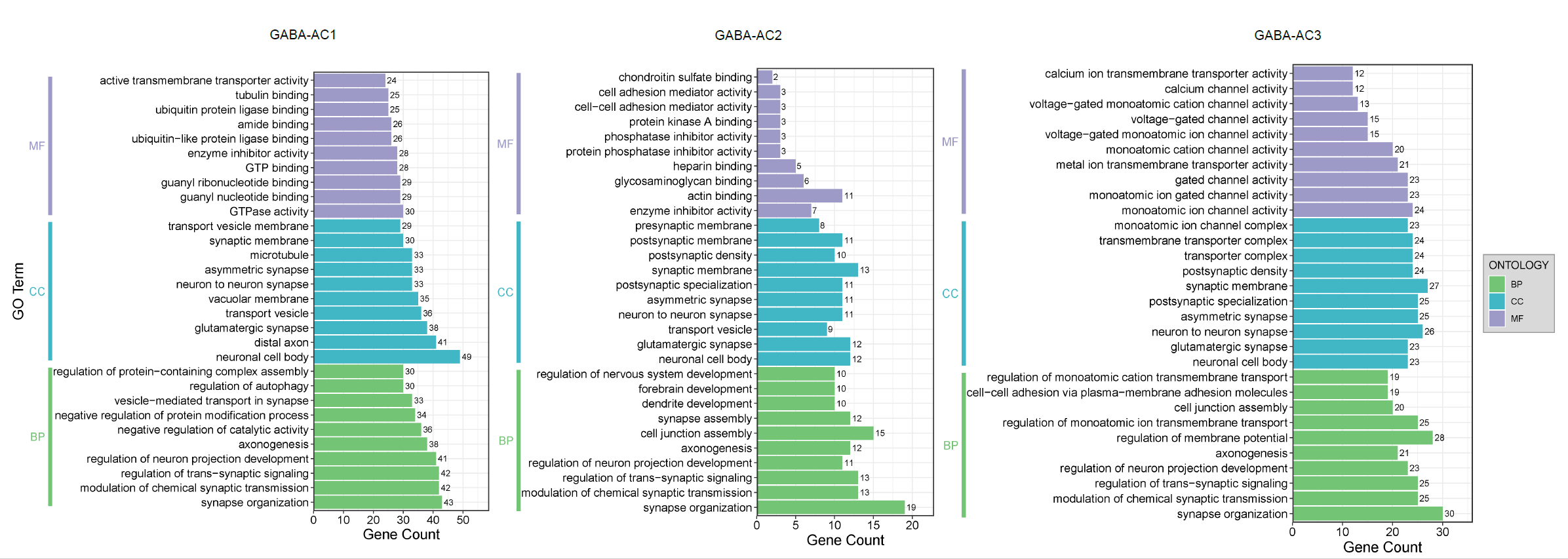

**FigS1 Gene Ontology (GO) analysis of the GABA-AC1, GABA-AC2, and GABA-AC3 subpopulations of amacrine cells.** In this figure, the top ten enriched GO terms are shown for each category, sorted by gene count. The bars represent the number of genes involved in each GO term, with the total gene count displayed on the right of each bar. The bar plots are divided into three sections based on Molecular Function (MF, purple), Cellular Component (CC, blue), and Biological Process (BP, green).

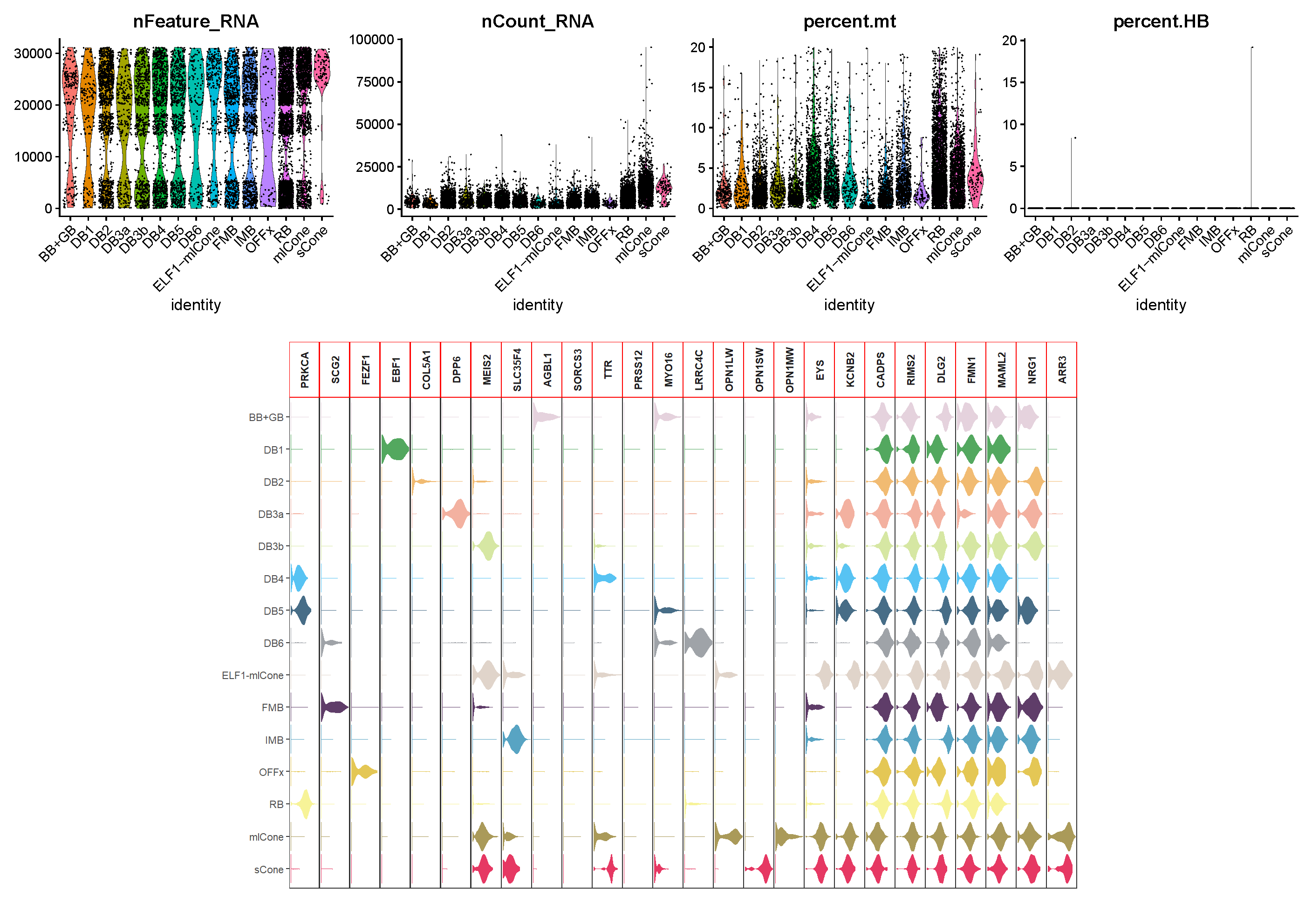

**FigS2** **Quality control and marker gene expression analysis of cone and bipolar cell subpopulations.**

**Top panel:** Violin plots showing quality control metrics for cone and bipolar cell subpopulations, including nFeature_RNA (number of detected genes), nCount_RNA (total RNA counts), percent.mt (percentage of mitochondrial RNA), and percent.HB (percentage of hemoglobin RNA). Each violin corresponds to a specific cell subpopulation.

**Bottom panel:** Violin plots showing the expression levels of key marker genes across cones and bipolar cell subpopulations. Each row corresponds to a subpopulation, and each column represents a distinct marker gene.

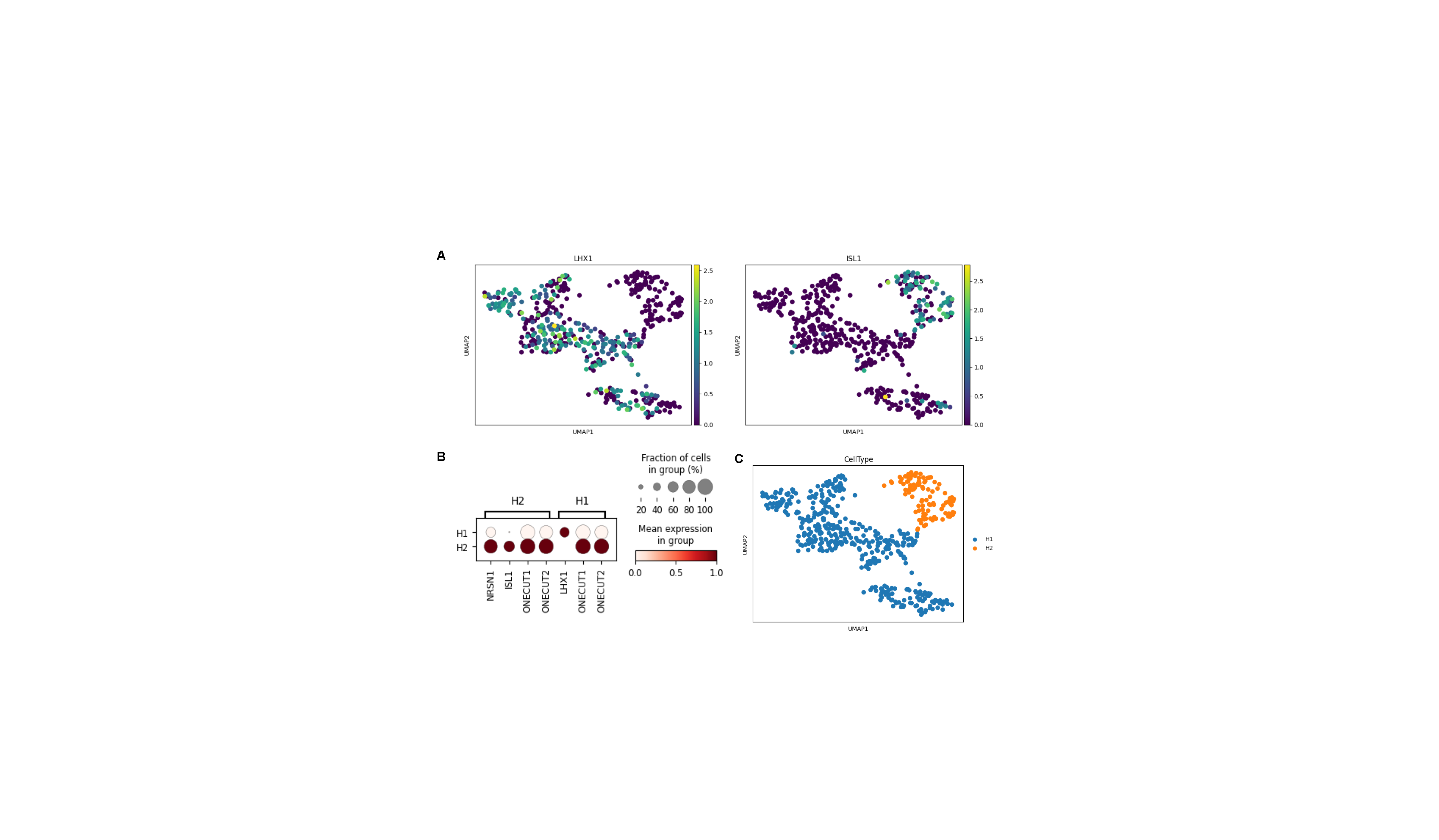

**FigS3 Analysis of gene expression in horizontal cell subtypes.**

(A) UMAP plots showing the expression of LHX1 and ISL1 across horizontal cell subtypes. The color scale represents the expression level of each gene.

(B) Dot plot summarizing key marker gene expression levels in H1 and H2 subtypes. Rows correspond to H1 and H2, and columns correspond to a list of key marker genes, brackets on the top detail the horizontal cell subtype that these genes identify.

(C) UMAP plot showing the clustering of horizontal cell subtypes based on gene expression profiles, with H1 cells in blue and H2 cells in orange.

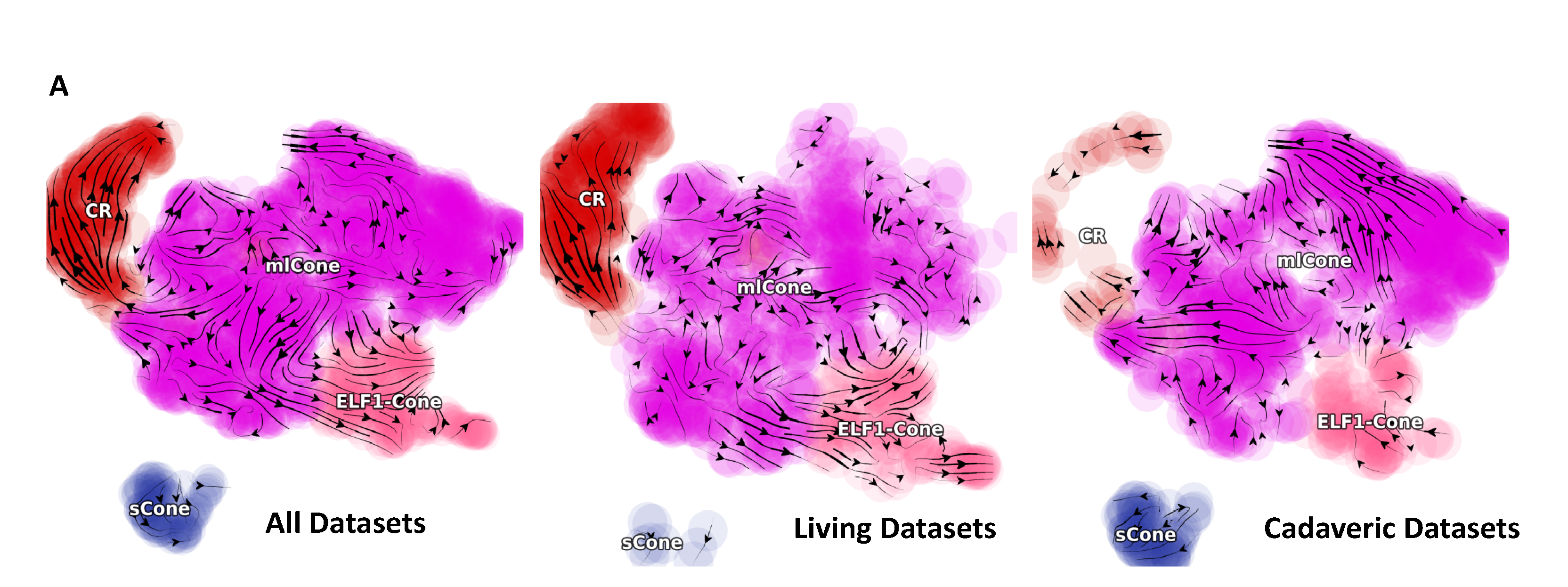

**FigS4 Dynamo analysis of ELF1-mlCone differentiation and regulation in human retina.** UMAP plots showing cell type distributions and velocity vectors across all datasets, living datasets, and cadaveric datasets. Cell types are colored by: CR cluster (red), mlCone (purple), ELF1-mlCone (pink), and sCone (blue). Arrows *represent* predicted cell state transitions.
